# Pan-genome and phylogeny of *Bacillus cereus sensu lato*

**DOI:** 10.1101/119420

**Authors:** Adam L. Bazinet

## Abstract

**Background:** *Bacillus cereus sensu lato* (*s. l*.) is an ecologically diverse bacterial group of medical and agricultural significance. In this study, I use publicly available genomes to characterize the *B. cereus s. l.* pan-genome and perform the largest phylogenetic and population genetic analyses of this group to date in terms of the number of genes and taxa included. With these fundamental data in hand, I identify genes associated with particular phenotypic traits (i.e., “pan-GWAS” analysis), and quantify the degree to which taxa sharing common attributes are phylogenetically clustered.

**Methods:** A rapid *k*-mer based approach (Mash) was used to create reduced representations of selected *Bacillus* genomes, and a fast distance-based phylogenetic analysis of this data (FastME) was performed to determine which species should be included in *B. cereus s. l.* The complete genomes of eight *B. cereus s. l.* species were annotated de novo with Prokka, and these annotations were used by Roary to produce the *B. cereus s. l.* pan-genome. Scoary was used to associate gene presence and absence patterns with various phenotypes. The orthologous protein sequence clusters produced by Roary were filtered and used to build HaMStR databases of gene models that were used in turn to construct phylogenetic data matrices. Phylogenetic analyses used RAxML, DendroPy, ClonalFrameML, PAUP*, and SplitsTree. Bayesian model-based population genetic analysis assigned taxa to clusters using hierBAPS. The genealogical sorting index was used to quantify the phylogenetic clustering of taxa sharing common attributes.

The *B. cereus s. l.* pan-genome currently consists of ≈60,000 genes, ≈600 of which are “core” (common to at least 99% of taxa sampled). Pan-GWAS analysis revealed genes associated with phenotypes such as isolation source, oxygen requirement, and ability to cause diseases such as anthrax or food poisoning. Extensive phylogenetic analyses using an unprecedented amount of data produced phylogenies that were largely concordant with each other and with previous studies. Phylogenetic support as measured by bootstrap probabilities increased markedly when all suitable pan-genome data was included in phylogenetic analyses, as opposed to when only core genes were used. Bayesian population genetic analysis recommended subdividing the three major clades of *B. cereus s. l.* into nine clusters. Taxa sharing common traits and species designations exhibited varying degrees of phylogenetic clustering.

## Background

*Bacillus cereus sensu lato* (*s. l.*) is an ecologically diverse bacterial group that comprises a growing number of species, many of which are medically or agriculturally important. Historically recognized and most well-sampled of the species are *B. anthracis* (the causative agent of anthrax), *B. cereus sensu stricto* (capable of causing food poisoning and other ailments), and *B. thuringiensis* (used to control insect pests). Other species are distinguished by rhizoidal growth patterns (*B. mycoides* and *B. pseudomycoides* [48]), thermotolerance and cytotoxicity (*B. cytotoxicus* [25]), psychrotolerance and ability to cause food spoilage (*B. weihenstephanensis* [38] and *B. wiedmannii* [46]), and utility as a probiotic in animal nutrition (*B. toyonensis* [30]). In addition, several new species have also recently been described (*B. bingmayongensis* [42], *B. gaemokensis* [32], and *B. manliponensis* [31]). In order to understand the fantastic diversity of *B. cereus s. l.* and its concomitant ability to occupy diverse environmental niches and exhibit a variety of phenotypes, it is crucial to accurately characterize genomic diversity within the group and to generate robust phylogenetic hypotheses about the evolutionary relationships among group members.

A typical *B. cereus s. l.* genome contains ≈5,500 protein-coding genes [53, 64]. Due to rampant horizontal gene transfer in bacterial ecosystems, however, the genome of a particular strain or species often contains genes not found in closely related taxa [63]. Thus, it is now common practice to seek to characterize the full gene complement of a closely related group of bacterial strains or species, otherwise known as a “pangenome” [63]. In this study, a “core” gene is defined as present in at least 99% of sampled taxa, an “accessory” gene as a non-core gene present in at least two taxa, and a “unique” gene as present in only one taxon. A previous effort to characterize the *B. cereus s. l.* pan-genome [37] based on a comparison of a relatively small number of strains estimated that there are ≈3,000 core genes and ≈22,500 total genes in the *B. cereus s. l.* pan-genome. A more recent study [69] using 58 strains reported similar estimates.

Phylogenetic hypotheses of *B. cereus s. l.* have been generated from a variety of data sources, including 16S rRNA sequences [37], amplified fragment length polymorphism (AFLP) data [26, 65], multilocus sequence typing (MLST) of housekeeping genes [8, 18, 20, 65], single-copy protein-coding genes [58], locally collinear blocks (LCBs) [69], conserved protein-coding genes [69], whole-genome single nucleotide polymorphisms (SNPs) [8], and digital DNA-DNA hybridization (dDDH) data [43]. Phylogenetic analyses have used distance methods [20, 26, 43, 69], maximum likelihood [8, 26, 58], maximum parsimony [26], and Bayesian methods [18]. For the most part, published phylogenies have tended to agree with and reinforce one another, although naturally there have been different classification systems developed with attendant implications for species designations. One popular classification system divides the *B. cereus s. l.* phylogeny into three broad clades [18,49,69]; traditionally, Clade 1 contains *B. anthracis, B. cereus*, and *B. thuringiensis;* Clade 2 contains *B. cereus* and *B. thuringiensis;* and Clade 3 contains a greater diversity of species including *B. cereus, B. cytotoxicus, B. mycoides, B. thuringiensis, B. toyonensis*, and *B. weihenstephanensis.* A somewhat more fine-grained classification system divides the phylogeny into seven major groups [8, 26, 65], each with its own thermotolerance profile [26] and propensity to cause food poisoning [27].

In this study I aimed to produce the most accurate and comprehensive estimate of the *B. cereus s. l.* pan-genome and phylogeny to date by analyzing all publicly available *B. cereus s. l.* genome data with a novel bioinformatic workflow for pan-genome characterization and pan-genome-based phylogenetic analysis.

## Methods

### Distance-based phylogeny of the genus *Bacillus*

All “reference” and “representative” *Bacillus* genome assemblies were retrieved from the NCBI RefSeq [50] database in October 2016, comprising 86 assemblies from 74 well-described *Bacillus* species and 44 assemblies from as-yet uncharacterized species. In addition, 16 “latest” assemblies were added for five *Bacillus* species that are thought to be part of *B. cereus s. l. (B. bingmayongensis* [42], *B. gaemokensis* [32], *B. pseudomycoides* [48], *B. toyonensis* [30], and *B. wiedmannii* [46]). In total, 146 *Bacillus* genomes were included in the distance-based phylogenetic analysis. The sketch function in Mash [51] version 1.1.1 (arguments: −k 21 −s 1000) was used to create a compressed representation of each genome, and then the Mash distance function was used to generate all pairwise distances among genomes. The Mash distance matrix was converted to PHYLIP format and analyzed with FastME [39] version 2.1.4 using the default BIONJ [24] algorithm.

### Creation of taxon sets

#### bcsl_114

All complete genomes of eight *B. cereus s. l.* species (*B. anthracis, B. cereus, B. cytotoxicus* [25], *B. mycoides, B. pseudomycoides* [48], *B. thuringiensis, B. toyonensis* [30], and *B. weihenstephanensis* [38]) were downloaded from the NCBI RefSeq [50] database in October 2016, which altogether comprised 114 genomes. One strain from each species was designated the “reference taxon” for that species, as required by HaMStR [21] (Table 1). This taxon set of complete genomes (“BCSL_114”; Table 1) was used to build the HaMStR databases and as the basis for the majority of the analyses performed in this study.

**Table 1.** Species composition of taxon sets.

| Species | Representatives<br>in BCSL_114 | Representatives<br>in BCSL_498 | HaMStR<br>reference taxon | RefSeq assembly<br>accession |
| --- | --- | --- | --- | --- |
| *B. anthracis* | 42 | 128 | Ames | GCF_000007845 |
| *B. bingmayongensis* | 0 | 1 |  |  |
| *B. cereus (sensu stricto)* | 30 | 258 | ATCC 14579 | GCF_000007825 |
| *B. cytotoxicus* | 1 | 2 | NVH 391-98 | GCF_000017425 |
| *B. gaemokensis* | 0 | 2 |  |  |
| *B. manliponensis* | 0 | 1 |  |  |
| *B. mycoides* | 2 | 13 | ATCC 6462 | GCF_000832605 |
| *B. pseudomycoides* | 1 | 1 | DSM 12442 | GCF_000161455 |
| *B. thuringiensis* | 35 | 73 | 97-27 | GCF_000008505 |
| *B. toyonensis* | 1 | 1 | BCT-7112 | GCF_000496285 |
| *B. weihenstephanensis* | 2 | 6 | KBAB4 | GCF_000018825 |
| *B. wiedmannii* | 0 | 11 |  |  |
| *Bacillus sp.* | 0 | 1 |  |  |

#### BCSL 498

To perform analyses involving all publicly available *B. cereus s. l.* genome data, all “latest assemblies” were downloaded for the eight species mentioned above, and based on analysis of the *Bacillus* distance-based phylogeny, assemblies were added for *B. bingmayongensis* [42], *B. gaemokensis* [32], *B. manliponensis* [31], *B. wiedmannii* [46], and one uncharacterized species (*Bacillus sp.* UNC437CL72CviS29), which altogether comprised 498 genomes (“BCSL 498”; Table 1). A list of RefSeq assembly accessions for all taxa used in this study is provided (Additional file 1).

### Isolate metadata

*B. cereus s. l.* isolate metadata, including “Assembly Accession”, “Disease”, “Host Name”, “Isolation Source”, “Motility”, and “Oxygen Requirement” was downloaded from PATRIC [67] in December 2016. This metadata was used to associate patterns of gene presence and absence with phenotypes exhibited by groups of taxa.

### Genome annotation

All *B. cereus s. l.* genomes were annotated de novo with Prokka [59] version 1.12-beta (arguments: `--kingdom Bacteria --genus Bacillus`).

### Pan-genome inference

The pan-genome of *B. cereus s. l.* was inferred with Roary [52] version 3.7.0. The bcsl_114 Prokka annotations were provided to Roary as input; in turn, Roary produced a gene presence/absence matrix (Additional file 2), a multi-FASTA alignment of core genes using PRANK [44] version 0.140603, and a tree based on the presence and absence of accessory genes among taxa using FastTree 2 [55] version 2.1.9. The “accessory binary tree” was computed using only the first 4,000 genes in the accessory genome.

### Phylogenetic network analysis

A NEXUS-format binary version of the bcsl_114 gene presence/absence matrix was analyzed with SplitsTree4 [28] version 4.14.4. Three methods of calculating distances between taxa were evaluated: `Uncorrected P, GeneContentDistance` [29], and the `MLDistance` variant of `GeneContentDistance` [29]. The `NeighborNet` [11] algorithm was used to reconstruct the phylogenetic network.

### Genotype-phenotype association

Scoary [12] version 1.6.9 was used to associate patterns of gene presence and absence with particular phenotypes (traits), an analysis known as “pan-GWAS” [12]. Scoary required two basic input files: the BCSL_114 gene presence/absence matrix, augmented with gene presence/absence information for bcsl_498 taxa obtained from orthology determination with HaMStR [21] (Additional file 3), and a binary trait matrix that was created using the isolate metadata obtained from PATRIC (Additional file 4). Assignment of traits to taxa was performed conservatively in that missing data was not assumed to be an indication of the presence or absence of a particular trait. Scoary was run with 1,000 permutation replicates, and genes were reported as significantly associated with a trait if they attained a naive *P*-value less than 0.05, a Benjamini-Hochberg-corrected *P*-value less than 0.05, an empirical *P*-value less than 0.05, and were not annotated as “hypothetical proteins”. Lists of genes were subsequently tested for enrichment of biological processes using the data and services provided by AmiGO 2 [13] version 2.4.24, which in turn used the PANTHER database [45] version 11.1.

### HaMStR database creation

The orthologous protein sequence clusters output by Roary were filtered to produce a set of gene models suitable for use with HaMStR [21] version 13.2.6. HaMStR enables one to build gene models for a clade of interest (using, ideally, high-quality complete genomes), which are subsequently used to identify orthologs in other sequence data (e.g., draft genome assemblies, transcriptomes, etc.). HaMStR required that each sequence cluster contain at least one sequence from the set of previously selected reference taxa (Table 1), so clusters not meeting this requirement were omitted. Furthermore, each cluster was required to contain at least four sequences (the minimum number of sequences required to produce an informative unrooted phylogenetic tree), and all cluster sequences needed to be at least 100 nt in length. Finally, clusters that Roary flagged as having a quality-control issue were removed. The 9,070 clusters that passed these filters were aligned using the linsi algorithm in MAFFT [35] version 7.305. Gene models (i.e., Hidden Markov Models, or HMMs) were produced from the aligned cluster sequences using the hmmbuild program from HMMER [22] version 3.0. Finally, for each reference taxon, a BLAST [1] database was built using the full complement of protein-coding genes for that taxon. This completed the construction of the initial HaMStR database, which is called “hamstr_full”. A variant of hamstr_full called “hamstr_core” was created, which contained only the 594 gene models corresponding to core genes.

#### Mobile genetic element removal

For tree-based phylogenetic analyses that assume a process of vertical inheritance, the inclusion of mobile genetic elements (MGEs) that may be horizontally transferred is likely to confound the phylogenetic inference process [9]. Thus, an effort was made to identify and remove putative MGEs from the HaMStR databases. In December 2016, a list of *Bacillus* genes derived from a plasmid source was downloaded from the NCBI Gene [15] database. In addition, all genes were exported from the ACLAME [40] database version 0.4. Using this information, gene models that were either plasmid-associated or found in the ACLAME list of MGEs were removed from HaMStR databases. Gene models whose annotation included the keywords “transposon”, “transposition”, “transposase”, “insertion”, “insertase”, “plasmid”, “prophage”, “intron”, “integrase”, or “conjugal” were also removed. The resulting HaMStR databases, “hamstr_full_mges_removed” and “hamstr_core_mges_removed”, contained 8,954 and 578 gene models, respectively. The workflow used to construct the hamstr_full_mges_removed database is shown as a diagram (Additional file 5).

### Orthology determination

The protein-coding gene annotations of “query” taxa — i.e., taxa not included in BCSL_114 — were searched for sequences matching HaMStR database gene models using HaMStR [21] version 13.2.6 (which in turn used GeneWise [6] version 2.4.1, HMMER [22] version 3.0, and BLASTP [1] version 2.2.25+). In the first step of the HaMStR search procedure, the `hmmsearch` program from HMMER was used to identify translated substrings of protein-coding sequence that matched a gene model in the database, which were then provisionally assigned to the corresponding sequence cluster. To reduce the number of highly divergent, potentially paralogous sequences returned by this initial search, the E-value cutoff for a “hit” was set to 1e–05 (the HaMStR default was 1.0). In the second HaMStR step, BLASTP was used to compare the hits from the HMM search against the proteome of the reference taxon associated with that gene model; sequences were only retained if the reference taxon protein used in the construction of the gene model was also the best BLAST hit. The E-value cutoff for the BLAST search was set to 1e-05 (the HaMStR default was 10.0).

### Data matrix construction

Amino acid sequences assigned to orthologous sequence clusters were aligned using MAFFT [35] version 7.305. The resulting amino acid alignments were converted to corresponding nucleotide alignments using a custom Perl script that substituted for each amino acid the proper codon from the original coding sequence. Initial orthology assignment may sometimes result in multiple sequences for a particular taxon/locus combination [4], which need to be reduced to a single sequence for inclusion in phylogenetic data matrices. For this task the “consensus” [3] procedure was used, which collapsed all sequence variants into a single sequence by replacing multi-state positions with nucleotide ambiguity codes. Following application of the consensus procedure, individual sequence cluster alignments were concatenated, adding gaps for missing data as necessary using a custom Perl script. The workflow used for orthology determination and data matrix construction is shown as a diagram (Additional file 6).

### Maximum likelihood phylogenetic analysis

Concatenated nucleotide data matrices were analyzed under the maximum likelihood criterion using RAxML [60] version 8.2.8 (arguments: `-f d -m GTRGAMMAI`). The data were analyzed either with all nucleotides included in a single data subset (all_nuc), or with sites partitioned by codon position (codon_pos). Partitioned analyses assigned a unique instance of the substitution model to each data subset, with joint branch length optimization. Analyses of the bcSl_114 taxon set consisted of an adaptive best tree search [5] and an adaptive bootstopping procedure that used the `autoMRE RAxML` bootstopping criterion [54]; thus, the number of search replicates performed varied from 10 to 1000, depending on the analysis. DendroPy [61] was used to map bootstrap probabilities onto the best tree. Analysis of the bcSl_498 taxon set required ≈256 GB of RAM and multiple weeks of runtime, and was thus limited to a single best tree search.

### Recombination detection

Genomic regions that may have been involved in past recombination events should be excluded from phylogenetic analyses that assume a process of vertical inheritance, or phylogenetic inference methods should incorporate this information to produce a more accurate phylogeny [9]. In this study, two different software packages that address this problem were evaluated. First, the `profile` program from PhiPack [10] was used to flag and remove from concatenated data matrices sites that exhibited signs of mosaicism. Following the procedure employed in Parsnp [66], the `profile` program defaults were used, except that the step size was increased from 25 to 100 (`-m 100`). RAxML was then used to create new versions of data matrices that excluded regions whose Phi statistic *P*-value was less than 0.01. Second, ClonalFrameML [19] (downloaded from GitHub June 14, 2016) with default parameters was used to correct the branch lengths of phylogenies to account for recombination. ClonalFrameML required all ambiguous bases in data matrices to be coded as “N”.

### Maximum parsimony phylogenetic analysis

Concatenated nucleotide data matrices were analyzed under the maximum parsimony criterion using PAUP* [62] version 4.0a150. A heuristic search was performed using default parameters.

### Tree distance calculation

To quantify the difference between pairs of tree topologies, both the standard and normalized Robinson-Foulds distance [57] were calculated with RAxML [60] version 8.2.8 (arguments: `-f r -z`).

### Tree visualization

Visualizations of phylogenetic trees were produced with FigTree [56] version 1.4.2, except Fig. 4, which was produced with iTOL [41] version 3.5.3.

### Taxon clustering

To complement phylogenetic analysis and existing classification systems, taxa were clustered with hierBAPS [14] (bugfixed version dated August 15, 2013), a Bayesian model-based population genetic approach that accounts for admixture within and among lineages. The bcSl_498 alignment of 8,954 genes (mat_6) was provided to hierBAPS as input, and hierBAPS was directed to produce a single-level clustering with a maximum of 10 clusters.

### Clustering of taxon-associated attributes

The degree of clustering of taxa sharing a common attribute, given a phylogeny relating those taxa, was quantified using the genealogical sorting index [16] (*gsi*) version 0.92 made available through the web service at `molecularevolution.org` [2]. Significance of the *gsi* was determined by running 10^4^ permutation replicates.

## Results

### Distance-based phylogeny of the genus *Bacillus*

The Mash-distance-based phylogeny of the genus *Bacillus* (Additional file 7) indicated a *B. cereus s. l.* clade containing the following species: *B. anthracis, B. bingmayongensis, B. cereus* (*sensu stricto), B. cytotoxicus, B. gaemokensis, B. manliponensis, B. mycoides, B. pseudomycoides, B. thuringiensis, B. toyonensis, B. weihenstephanensis, B. wiedmannii*, and one uncharacterized species (*Bacillus sp.* UNC437CL72CviS29). Within *B. cereus s. l.*, the first taxon to split off from the remainder of the group was *B. manliponensis*, followed by *B. cytotoxicus* (which has been previously recognized as an outlier [37,58]).

### Pan-genome inference

The pan-genome of *B. cereus s. l.* was inferred with Roary [52] using the bcsl_114 taxon set. Roary produced a total of 59,989 protein-coding gene sequence clusters (Additional file 2) from an average of 5,726 genes per input genome (Additional file 8). The shortest cluster sequence was 122 nt, the longest cluster sequence was 22,967 nt, and the average length of a cluster sequence was 755 nt. The average difference between the shortest and longest sequence in a cluster was only 67 nt, suggesting that the input data was relatively uniform, as would be expected from complete genomes. The *B. cereus s. l.* “core genome”, consisting of genes present in at least 99% of taxa sampled, was represented by 598 genes (≈1% of all genes). A rarefaction curve shows that after ≈35 genomes have been sampled (≈31% of all genomes), the number of core genes remains fairly constant at ≈600 genes, while the total number of genes in the pan-genome continues to increase almost linearly (Additional file 10). The 59,391 non-core genes were divided into 32,324 “accessory genes” (i.e., non-core genes present in at least two taxa; ≈54% of all genes), and 27,067 “unique genes” (i.e., genes present in only one taxon; ≈45% of all genes). A rarefaction curve shows that as genomes are sampled, genes never before observed continue to be found at a fairly steady rate, and the total number of unique genes discovered continues to increase, with no indication of soon approaching an asymptote (Additional file 11); these trends indicate that the *B. cereus s. l.* pan-genome is “open”. Finally, Roary produced an “accessory binary tree”, which was plotted alongside core and accessory gene presence/absence information (Additional file 12). This figure shows that the outermost *B. cereus s. l.* clades include taxa with relatively few accessory genes included in the analysis (≤ 40), such as *B. cytotoxicus, B. mycoides*, and *B. pseudomycoides;* by contrast, the genomes with the most accessory genes present (> 1000) belong to the highly clonal clade of *B. anthracis* strains.

### Phylogenetic network analysis

SplitsTree4 [28] was used to build a phylogenetic network from the gene presence/absence information produced by Roary. The choice of method for computing distance did affect network branch lengths; the network presented here was computed with the `MLDistance` variant of `GeneContentDistance` (Fig. 1), as that method seemed most appropriate for gene presence/absence data. The phylogenetic network recapitulated both the three-clade [18, 49, 69] and seven-group [8, 26, 65] classification systems used in previous studies. Group I and Group VII, both part of Clade 3, were most radically diverged from the remainder of the network. Notably, Group II was absent from the network, as it was not represented by any complete *B. cereus s. l.* genomes at the time the study was performed.

**Figure 1.**
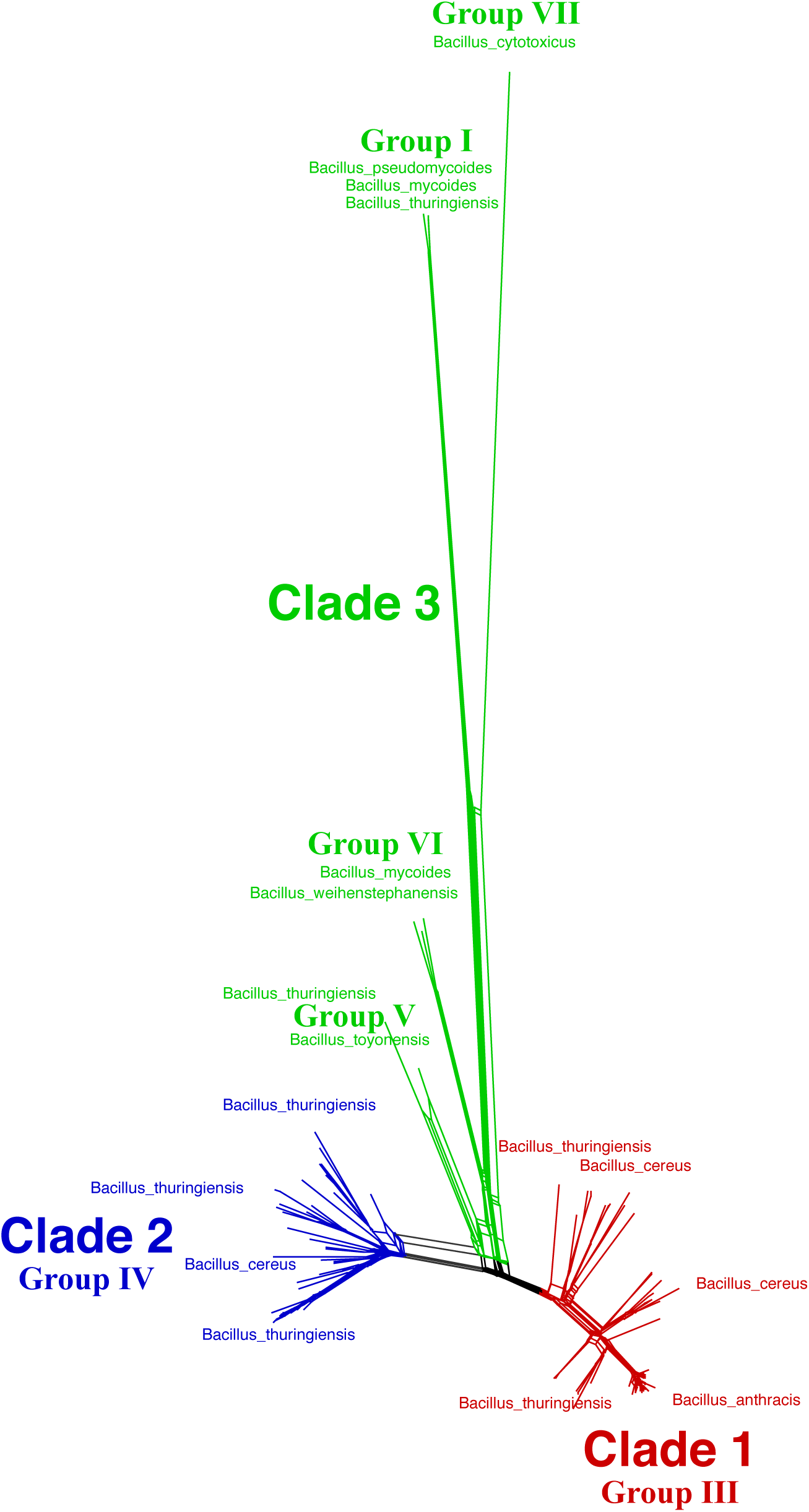
Phylogenetic network analysis of BCSL_114. Gene presence/absence information produced by Roary was provided as input to SplitsTree, which used the `MLDistance` variant of `GeneContentDistance` together with the NeighborNet algorithm to reconstruct the phylogenetic network. Major *B. cereus s. l.* clades and groups are indicated, along with representative taxa.

### Genotype-phenotype association

Scoary [12] was used to associate patterns of gene presence and absence with particular phenotypes (traits), an analysis known as “pan-GWAS” [12]. Pan-GWAS was performed for the following traits: isolation source (cattle, human, invertebrate, non-primate mammal, or soil); motility; oxygen requirement (aerobic or facultative); and disease (anthrax or food poisoning). Eight of ten traits tested had some number of significant positively or negatively associated genes (Table 2). Traits with a sufficient number of associated genes were tested for possible enrichment of gene ontology biological processes (Additional file 13). The most interesting findings from this analysis concerned taxa isolated from soil. Specifically, metabolic and biosynthetic processes involving quinone (and in particular, menaquinone) were positively associated with soil isolates. Analysis of quinone species present in soil have been used previously to characterize soil microbiota [23]. Furthermore, a high ratio of menaquinone to ubiquinone (the two dominant forms of quinone in soil) has been associated with the presence of gram-positive bacteria such as *Bacillus* species [34]. On the other hand, biological processes involving flagella, cilia, or motility more generally were negatively associated with soil isolates. This finding is consistent with observations that motility may not be necessary for bacterial colonization of plant roots [17], doubts about the evolutionary advantage of maintaining flagella in a soil environment [33], and general properties of soil that bring into question the importance of active movement and the extent to which it occurs [47].

**Table 2.** Scoary result summary.

| Trait | Taxa *with* trait | Taxa *without* trait | Significant<br>*positively*<br>associated genes | Significant<br>*negatively*<br>associated genes |
| --- | --- | --- | --- | --- |
| isolation source: cattle | 25 | 331 | 358 | 227 |
| isolation source: human | 44 | 312 | 53 | 47 |
| isolation source: invertebrate | 16 | 340 | 0 | 0 |
| isolation source: non-primate mammal | 46 | 310 | 162 | 85 |
| isolation source: soil | 121 | 235 | 34 | 34 |
| motility | 88 | 18 | 0 | 0 |
| oxygen requirement: aerobic | 65 | 41 | 15 | 18 |
| oxygen requirement: facultative | 41 | 65 | 20 | 11 |
| disease: anthrax | 85 | 63 | 44 | 1 |
| disease: food poisoning | 43 | 105 | 3 | 23 |

### Concatenated data matrices

In total, six different concatenated nucleotide data matrices were constructed and analyzed (mat_1–mat_6; Table 3). The majority of the data matrices used the bcsl_114 taxon set (mat_1–mat_5); only MAT 6 used the BCSL 498 taxon set. Various gene sets were used, including 1) all core genes identified by Roary (all_core); 2) only the core genes used to build the HaMStR database (hamstr_core); 3) HaMStR core genes with mobile genetic elements (MGEs) removed (hamstr_core_mges_removed), and a variant of this gene set with PhiPack sites removed; and finally, 4) all HaMStR genes with MGEs removed (hamstr_full_mges_removed). Aligned data matrices ranged from 96,802 nt to 8,207,628 nt in length. Matrix completeness, defined as the percentage of non-missing data, ranged from 47.4% to 99.5%. The percentage of ambiguous characters present in data matrices ranged from 0.0% to 17.0%.

**Table 3.**
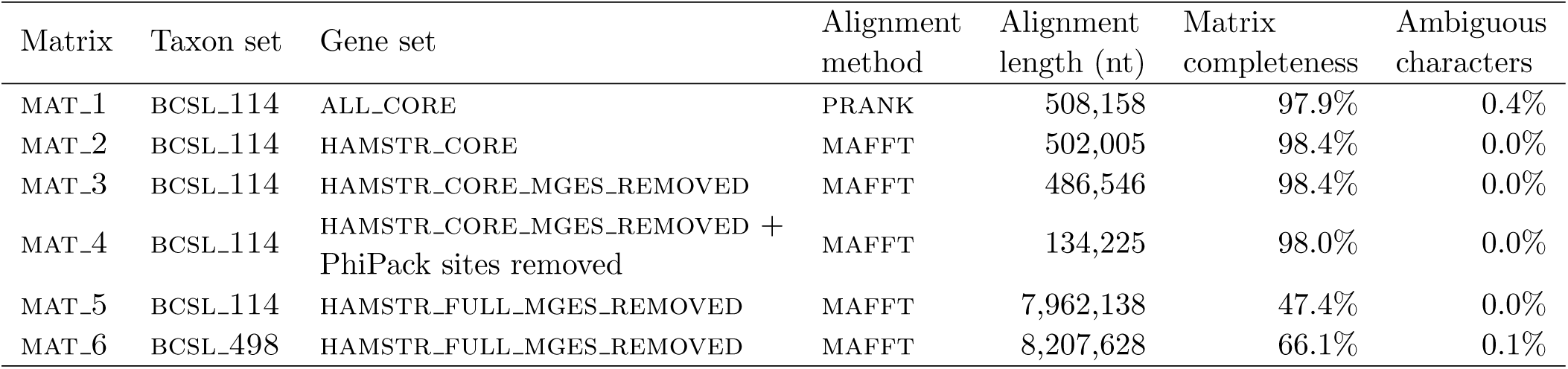
Concatenated data matrix statistics.

### Phylogenetic analyses

In total, nine different phylogenetic analyses of the six concatenated data matrices were performed (Table 4). Eight of the nine analyses used maximum likelihood (ml_1–ml_8), and one analysis used maximum parsimony (mp_1). For reasons of computational tractability, all exploratory analyses used the bcsl_114 taxon set (ml_1–ml_7 and mp_1); only when the best-performing methods were established was analysis of the bcsl_498 taxon set pursued (ml_8). During the exploratory phase, several variables were tested for their effect on phylogenetic outcome: 1) use of MAFFT instead of PRANK to align protein sequence clusters; 2) removal of MGEs; 3) use of maximum parsimony in addition to maximum likelihood; 4) partitioning of sites by codon position; 5) removal of sites implicated in recombination; and finally, 6) use of *all* eligible genes from the pan-genome versus only core genes.

**Table 4.** Phylogenetic analysis statistics^1^.

| Analysis | Matrix | Partitioning scheme | Unique alignment patterns | Best tree searches | Bootstrap replicates | Nodes with $BP = 1.0$ | Nodes with $BP \geq 0.5$ | Nodes with $BP < 0.5$ |
| --- | --- | --- | --- | --- | --- | --- | --- | --- |
| ML_1 | MAT_1 | ALL_NUC | 46,395 | 1000 | 100 | 68 | 98 | 14 |
| ML_2 | MAT_2 | ALL_NUC | 46,174 | 1000 | 100 | 68 | 93 | 19 |
| ML_3 | MAT_3 | ALL_NUC | 44,750 | 1000 | 200 | 63 | 94 | 18 |
| ML_4 | MAT_3 | CODON_POS | 49,889 | 1000 | 200 | 64 | 92 | 20 |
| ML_5 | MAT_4 | ALL_NUC | 11,729 | 1000 | 200 | 53 | 86 | 26 |
| ML_6 | MAT_6 | ALL_NUC | 691,147 | 10 | 100 | 92 | 112 | 0 |
| ML_7 | MAT_6 | CODON_POS | 852,707 | 28 | 100 | 89 | 112 | 0 |
| ML_8 | MAT_7 | ALL_NUC | 3,948,459 | 1 | 0 | N/A | N/A | N/A |
| MP_1 | MAT_3 | ALL_NUC | 83,383<sup>2</sup> | N/A | N/A | N/A | N/A | N/A |

<sup>1</sup> ML = maximum likelihood; MP = maximum parsimony; $BP$ = bootstrap probability; N/A = not applicable.
<sup>2</sup> Number of parsimony-informative characters.

Importantly, all phylogenetic analysis results recapitulated the three-clade [18, 49, 69] and seven-group [8, 26, 65] classification systems of previous studies. Taxa were consistently assigned to the same clade and group, independent of the particular phylogenetic analysis performed. Thus, topological differences between analysis results, as measured by the Robinson-Foulds distance [57] (Additional file 14), were confined to intra-group relationships. Bootstrap support was fairly consistent for all analyses that used core genes, and increased dramatically when all eligible genes from the pan-genome were used (Table 4). Additional detail about the phylogenetic analyses, and the logic behind their progression, is provided in the subsections that follow.

#### Choice of multiple sequence alignment program

Roary produced multiple sequence alignments of all 598 core genes with PRANK [44], which explicitly models insertions and deletions, but as a consequence runs more slowly than some other alignment programs. The PRANK alignments were concatenated to produce mat_1. A similar matrix was built using the 594 HaMStR-eligible core genes, except that the gene sequence clusters were aligned with MAFFT [35] (mat_2). Phylogenetic analyses of these two matrices with RAxML [60] revealed only negligible differences in bootstrap probabilities (ml_1 vs. ml_2; Table 4), so for the sake of computational efficiency MAFFT was used for the remainder of the analyses.

#### Removal of mobile genetic elements

For tree-based phylogenetic analyses that assume a process of vertical inheritance, the inclusion of mobile genetic elements (MGEs) that may be horizontally transferred is likely to confound the phylogenetic inference process [9]. Thus, putative MGEs were identified and removed from hamstr_core, leaving 578 core genes (hamstr_core_mges_removed). Phylogenetic analysis of this slightly smaller data matrix (mat_3) revealed comparable bootstrap probabilities to those from the analysis that used hamstr_core (ml_3 vs. ml_2; Table 4); nevertheless, out of principle, HaMStR databases with MGEs removed were used for the remainder of the analyses.

#### Partitioning of sites by codon position

It is well known that nucleotides in different codon positions (first, second, or third) are likely to be under different selective pressures [7]; thus, when analyzing protein-coding nucleotide sequences, it is common practice to apply a different substitution model (or different instance of the same substitution model) to the sites associated with each codon position, thus effectively partitioning the data matrix into three data subsets. The effect of partitioning by codon position was tested with two different matrices (mat_3 and mat_5); only negligible differences in bootstrap probabilities were found as compared to the unpartitioned results (ml_4 vs. ml_3 and ml_7 vs. ml_6; Table 4).

#### Removal of sites implicated in recombination

Genomic regions that may have been involved in past recombination events should not be used for phylogenetic analyses that assume a process of vertical inheritance [9]. The `profile` program from PhiPack [10] was used to flag and remove sites from mat_3 that exhibited signs of mosaicism. The resulting data matrix (mat_4) contained less than one-fourth the number of unique alignment patterns of mat_3, thus representing a substantial reduction in data suitable for phylogenetic analysis. This was reflected in bootstrap probabilities, which were somewhat depressed overall (ml_5 vs. ml_3; Table 4). It was thus concluded that removing sites implicated in recombination had a deleterious effect on phylogenetic analysis results, and so this procedure was not applied to subsequent analyses.

#### Use of all eligible genes from the pan-genome versus only core genes

Using all eligible genes (hamstr_full_mges_removed) for phylogenetic analysis as opposed to using only core genes (hamstr_core_mges_removed) caused bootstrap probabilities to increase dramatically (ml_6 vs. ml_3 and ml_7 vs. ml_4; Table 4). Thus, the ml_6 result was selected as the best estimate of the phylogenetic relationships among the bcsl_114 taxa. ClonalFrameML [19] was used to correct the branch lengths of this tree to account for recombination, and the tree was rooted using *B. cytotoxicus* [37, 58]. The resulting bcsl_114 phylogeny is shown as a phylogram with major clades and groups indicated (Fig. 2), and as a cladogram with bootstrap probabilities annotated (Additional file 15).

**Figure 2.**
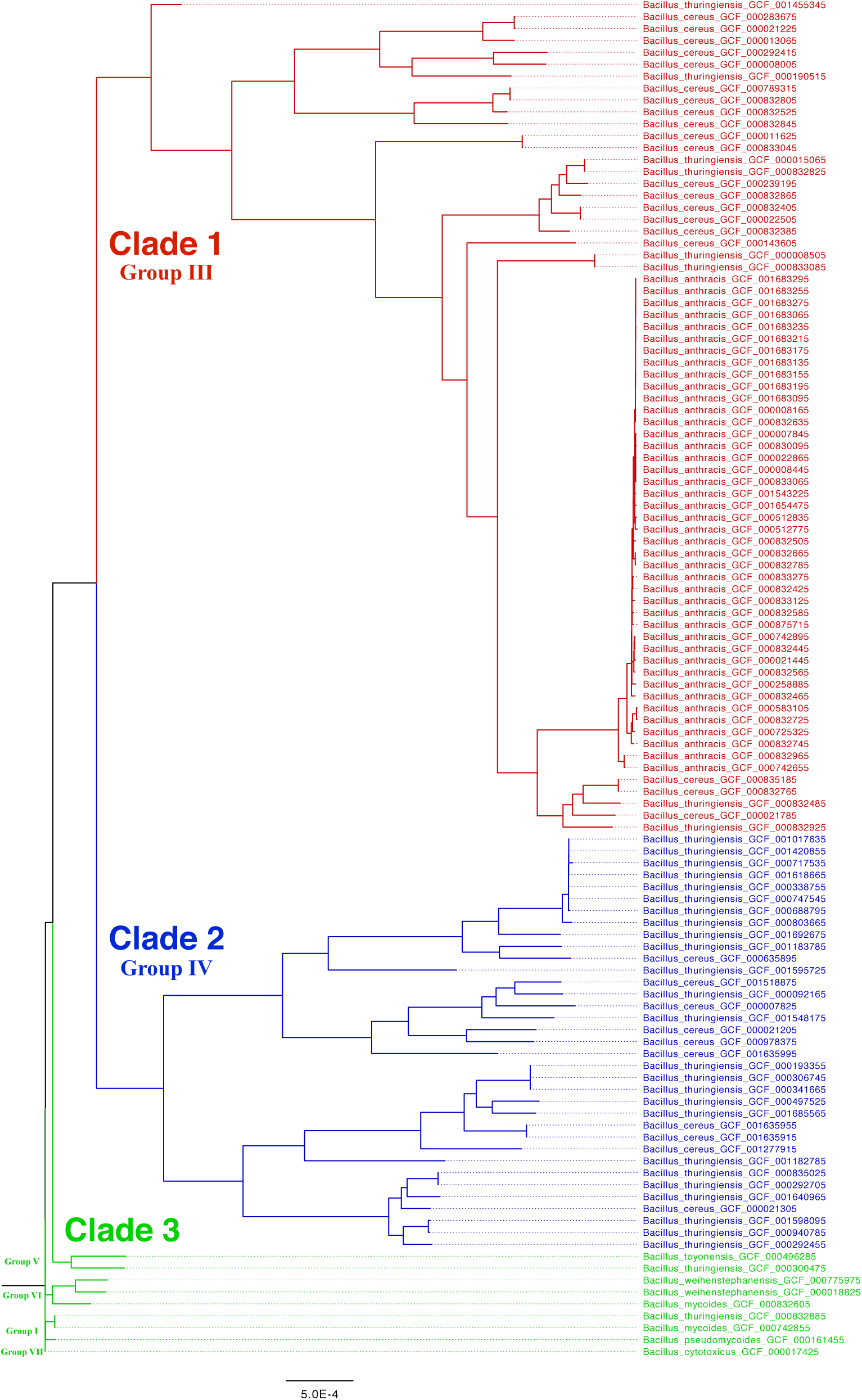
BCSL_114 maximum likelihood phylogenetic analysis results. Phylogram depicting the best estimate of the phylogenetic relationships among bcsl_114 taxa, computed with RAxML using 8,954 genes (ml_7; Table 4). ClonalFrameML was used to correct the branch lengths of the tree to account for recombination, and *B. cytotoxicus* was used to root the tree. Major *B. cereus s. l.* clades and groups are indicated.

#### Maximum likelihood-based analysis of all taxa

Once the exploratory analyses were completed, an analysis of bcsl_498 was executed using the hamstr_full_mges_removed gene set. The average number of genes included in the analysis for each species, clade, and group is given in Additional file 8, and a count of the number of genes included for each taxon is given in Additional file 9. Due to the size of the data matrix (almost 4 × 10^6^ unique alignment patterns), only a single best tree search replicate was completed (ml_8; Table 4). Informed by the distance-based analysis of *Bacillus* species (Additional file 7), the tree was rooted using *B. manliponensis.* The resulting bcsl_498 phylogeny is shown as a phylogram with major clades and groups indicated (Fig. 3). In contrast to analyses of bcsl_114, Group II is now represented, and is located on the tree where expected [8, 26, 65]. Based on this topology of currently sequenced genomes, a count of the number of taxa by species is provided for major *B. cereus s. l.* clades and groups (Additional file 8).

**Figure 3.**
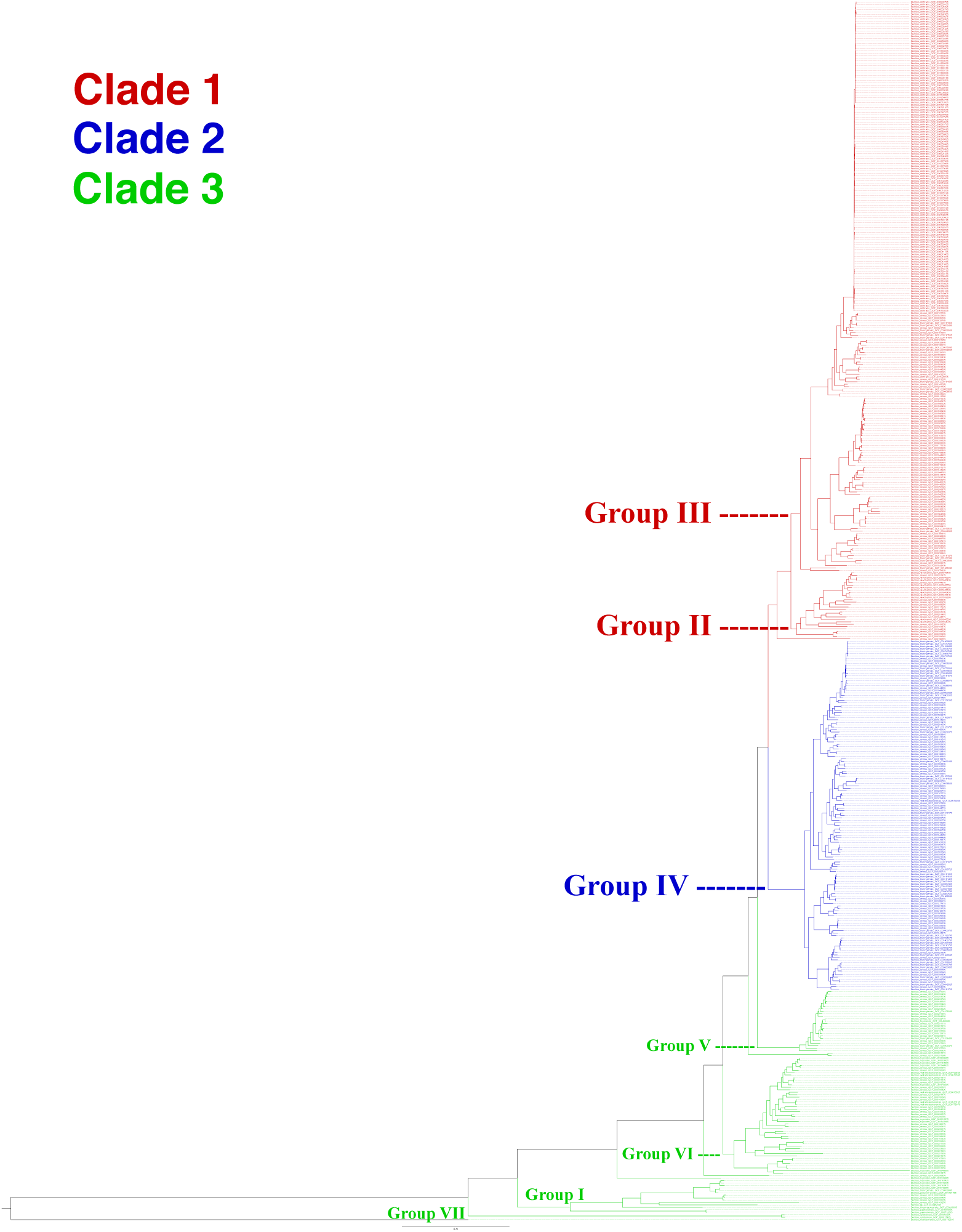
BCSL_498 maximum likelihood phylogenetic analysis results. Phylogram depicting an estimate of the phylogenetic relationships among bcsl_498 taxa, computed with RAxML using 8,954 genes (ml_9; Table 4). *B. manliponensis* was used to root the tree. Major *B. cereus s. l.* clades and groups are indicated.

### Taxon clustering

The hierBAPS [14] clustering analysis divided bcsl_498 into nine clusters (Additional file 9), which are displayed alongside major *B. cereus s. l.* clades and groups in Fig. 4. The hierBAPS clusters are congruent with the three-clade classification system, and largely agree with the seven-group classification system, with the following differences. Clade 1 included members of three clusters (as opposed to only two groups), and Clade 3 included members of six clusters (as opposed to four groups). Some Clade 3 clusters expanded slightly relative to their counterpart group to include taxa that were not assigned to any group, and *B. manliponensis* was assigned to its own cluster. Interestingly, two Clade 3 *B. cereus* taxa were assigned to Cluster 3, whereas other members of Cluster 3 were assigned to Clade 1, thus suggesting some genetic admixture between these two clades that was not reflected in the phylogenetic analysis.

**Figure 4.**
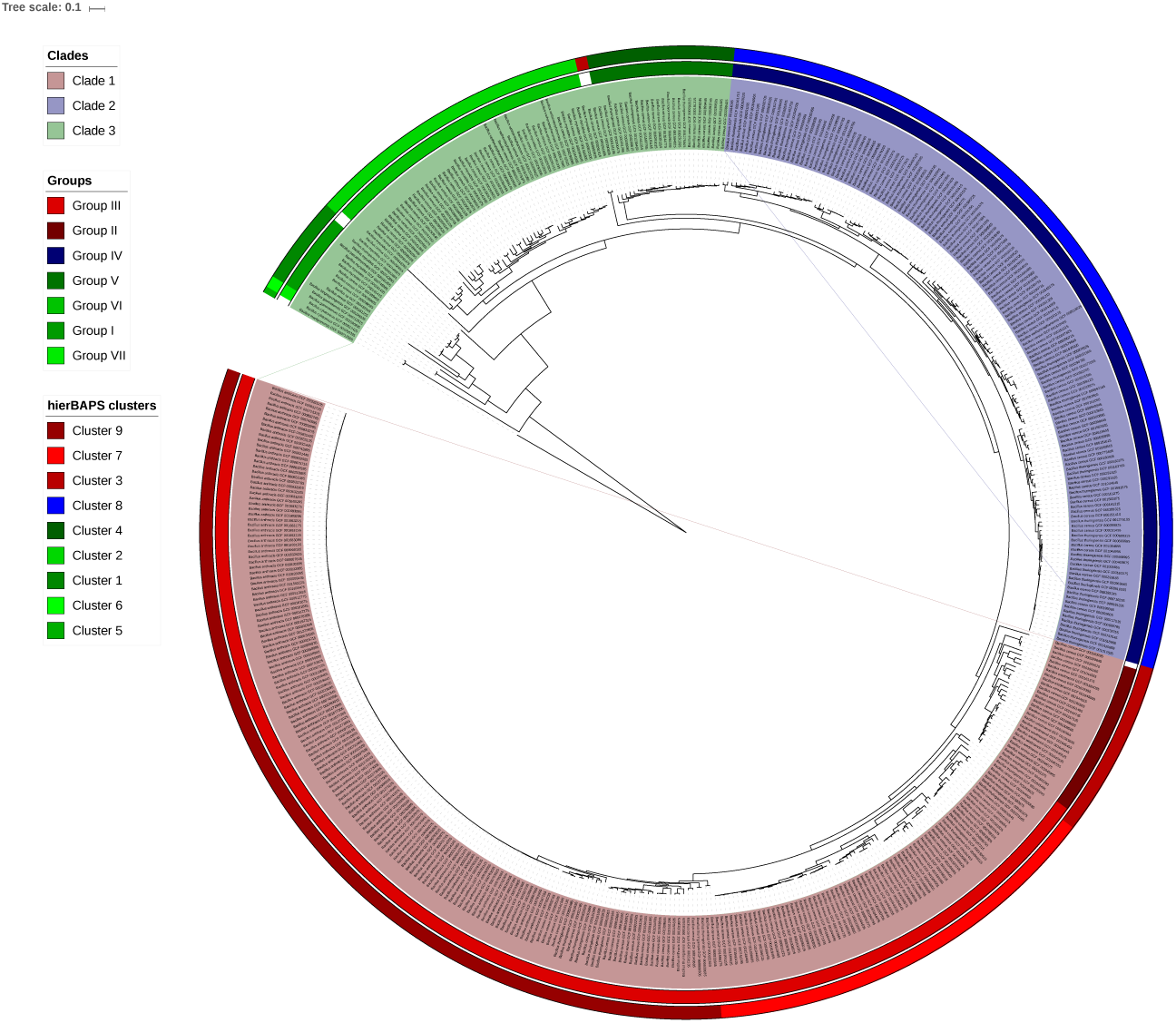
Phylogeny showing assignment of taxa to clades, groups, and clusters. Circular phylogram depicting an estimate of the phylogenetic relationships among bcsl_498 taxa, computed with RAxML using 8,954 genes (ml_8; Table 4). *B. manliponensis* was used to root the tree. Major *B. cereus s. l.* clades are indicated by highlighting of taxon labels, major groups are indicated by the inner colored strip, and hierBAPS clusters are indicated by the outer colored strip.

### Clustering of taxon-associated traits

The genealogical sorting index [16] (*gsi*) was used to quantify the degree of clustering of taxa sharing a common attribute given a phylogeny relating those taxa. The *gsi* statistic for a particular attribute takes a value from the unit interval [0,1]; if taxa associated with the attribute form a monophyletic group, the *gsi* = 1; otherwise, the greater the degree to which taxa associated with the attribute are dispersed throughout the tree (accounting for the number of taxa and the size of the tree), the smaller the *gsi* will be for that attribute.

#### Quantifying the degree of *B. cereus s. l.* species monophyly

The *gsi* was calculated for six *B. cereus s. l.* species that were sufficiently represented in the bcsl_498 phylogeny; all *P*-values were ≪ 0.05 (Additional file 16 and Table 5). Due to its highly clonal nature, *B. anthracis* was the species closest to monophyly (*gsi* = 0.95), and would have indeed been monophyletic except that one *B. anthracis* taxon (GCF 001029875) did not group with the others (but still placed in Group III). This might be a misannotation and should be investigated. *B. weihenstephanensis* was the species furthest from monophyly (*gsi* = 0.15), primarily because it was represented by only six taxa, one of which (GCF 000518025) was found in Group IV — the remainder were found in Group VI. Again, the annotation of the Group IV taxon with regard to species affiliation should be scrutinized.

**Table 5.** Monophyly status of *B. cereus s. l.* species, as quantified by the gsi.

| Species | Representatives in BCSL_498 | *gsi* value | *P*-value |
| --- | --- | --- | --- |
| *B. anthracis* | 128 | 0.95 | 0.0001 |
| *B. cereus* | 258 | 0.55 | 0.0001 |
| *B. mycoides* | 13 | 0.34 | 0.0001 |
| *B. thuringiensis* | 73 | 0.36 | 0.0001 |
| *B. weihenstephanensis* | 6 | 0.15 | 0.0021 |
| *B. wiedmannii* | 11 | 0.58 | 0.0001 |

#### Quantifying the degree of clustering of taxa sharing common traits

The *gsi* was calculated for ten traits shared by various *B. cereus s. l.* taxa using the bcsl_114 phylogeny from the ml_6 analysis; all *P*-values with the exception of one were less than 0.05 (Table 6). As not all of the taxa in bcsl_114 were assayed for each trait, the *gsi* values are artificially depressed; nevertheless, their relative values may be compared. The traits with the largest *gsi* values were “isolation source: cattle” and “isolation source: non-primate mammal”, the taxa associated with the former being a subset of the taxa associated with the latter. These taxa were all located in Group III, and all but two were identified as *B. anthracis.* This finding is consistent with the prevalence of mortality due to anthrax among cattle and other herbivores [68].

**Table 6.** Degree of clustering of taxa sharing common traits, as quantified by the gsi.

| Trait | Taxa with trait | *gsi* value | *P*-value |
| --- | --- | --- | --- |
| isolation source: cattle | 9 | 0.33 | 0.0001 |
| isolation source: human | 17 | 0.19 | 0.0273 |
| isolation source: invertebrate | 5 | 0.16 | 0.0462 |
| isolation source: non-primate mammal | 15 | 0.37 | 0.0001 |
| isolation source: soil | 17 | 0.27 | 0.0002 |
| motility | 25 | 0.16 | 0.2100 |
| oxygen requirement: aerobic | 15 | 0.19 | 0.0268 |
| oxygen requirement: facultative | 9 | 0.22 | 0.0037 |
| disease: anthrax | 11 | 0.26 | 0.0008 |
| disease: food poisoning | 8 | 0.17 | 0.0302 |

## Discussion

I show that the *B. cereus s. l.* pan-genome is still “open” (Additional files 10 and 11), thus implying that continued sampling of the group — especially of underrepresented taxa such as environmental strains [20] — will continue to reveal novel gene content. My estimate of the number of protein-coding genes in the *B. cereus s. l.* core and pan-genome (≈600 and ≈60,000, respectively), based on 114 complete genomes, is consistent with previous estimates [37, 69], as more extensive sampling of an open pan-genome will necessarily reduce the core genome size while simultaneously increasing the pan-genome size. It is interesting to observe that the basic phylogenetic structure of *B. cereus s. l.* can be accurately computed by relatively quick phylogenetic analyses based solely on the distribution of accessory genes among taxa (Fig. 1 and Additional file 12), which may in fact be sufficient for some applications. The diversity and adaptability of *B. cereus s. l.* may be in part attributable to the significant proportion of unique genes in its pan-genome (≈27,000, almost 50% of all genes; Additional file 11).

Pan-GWAS analysis found a number of genes significantly associated with various phenotypic traits (Table 2). In terms of validating this analysis, one might naturally look for genes known to be associated with *B. anthracis* virulence [36] or *B. cereus s. l.-* induced food poisoning [27]; however, these genes are not found among the analysis results. Many of these genes were not annotated by Roary, and of the ones that were, some were not represented in the hamstr_full database, thus reducing the number of taxa for which there would have been usable data. The genes that were reported to be significantly associated with “disease: anthrax”, “disease: food poisoning”, and other traits thus represent hypotheses that remain to be validated. Only four traits had enough significant positively or negatively associated genes to allow for the identification of enriched subsets of genes involved in particular biological processes (“isolation source: cattle”, “isolation source: human”, “isolation source: non-primate mammal”, and “isolation source: soil”; Additional file 13). Of these, only the biological processes associated with “isolation source: soil” were sufficiently specific so as to be meaningfully interpretable. To increase the statistical power of the pan-GWAS analysis and thereby generate more comprehensive and specific lists of genes associated with various traits, one would need to include additional taxa with relevant metadata and gene content information.

All phylogenetic analyses in this study recapitulated the three-clade and seven-group classification systems, and taxa were consistently assigned to the same clade and group (Figs. 1-4), irrespective of the data source or analysis methodology used (Tables 3 and 4). This strongly suggests that the broad phylogenetic structure of *B. cereus s. l.* has been inferred correctly. I demonstrate that the three-clade and seven-group systems are compatible with each other, as no group has its member taxa assigned to multiple clades. Clades 1 and 2 are much more extensively sampled than Clade 3 due to historical interest in *B. anthracis* and *B. thuringiensis* (Additional file 8); a recent study has shown that there is likely to be a tremendous amount of as-yet incompletely characterized diversity in Clade 3 that can be assayed by sampling various natural environments [20]. Indeed, Clade 3 exhibited the greatest degree of species diversity; in particular, Group I contained representatives of seven different species, including two newly characterized species (*B. bingmayongensis* [42] and *B. gaemokensis* [32]; Additional file 8). Six of the 498 taxa did not place into one of the seven previously circumscribed groups, which suggests that classification systems will need to be updated and refined as additional isolates are sequenced. Perhaps most interesting among the unplaced taxa is *B. manliponensis* [31], which appears to be even more divergent from other *B. cereus s. l.* taxa than *B. cytotoxicus* [25] (Fig. 3 and Additional file 7). One possibility for updating the group-level classification system is to incorporate information from the Bayesian model-based clustering analysis, the results of which were shown to be compatible with the three-clade system and which recommended nine clusters instead of seven groups (Fig. 4).

Using the phylogeny of bcsl_498, I quantified the degree of monophyly for six current *B. cereus s. l.* species designations (Additional file 16 and Table 5). This analysis demonstrates quantitatively that with the exception of *B. anthracis*, species definitions within *B. cereus s. l.* are not currently based on phylogenetic relatedness, but rather on phenotypes such as virulence, physiology, and morphology [8,26]. The primary focus of this study is the accurate reconstruction of phylogenetic relationships among taxa, and thus I make no specific recommendations for species re-designation based on these results. However, I do note a trend towards refined species designations that correlate with group affiliation; for example, several *B. cereus* strains in Group II have recently been re-designated *B. wiedmanii* [46]; similarly, Böhm et al. [8] suggested that all Group V taxa should be designated *B. toyonensis* [30]. In general, I recommend that taxonomic revisions are informed by well-supported phylogenetic hypotheses that have been generated without bias towards any particular species concept (e.g., dDDH boundaries [43]).

In a bioforensic setting, phylogenies that include well-supported strain-level relationships aid greatly in the identification of new isolates, and thus support both the attribution process (traceback of an isolate to its source) as well as analyses of pathogen evolution in an epidemic or outbreak scenario. However, the extremely high level of genomic conservation among closely related bacterial strains, especially in the core genome or in commonly typed conserved regions such as housekeeping genes, has limited the ability of previous analyses to make robust strain-level phylogenetic inferences. An important contribution of the current study is to show that bootstrap probabilities increase substantially when accessory genes are included in phylogenetic analyses along with core genes (Table 4). Thus, I have been able to resolve many strain-level, intra-group relationships of *B. cereus s. l.* with 100% bootstrap support for the first time (Additional file 15).

## Conclusion

In this study, I used novel bioinformatic workflows to characterize the pan-genome and phylogeny of *B. cereus sensu lato.* Based on data from 114 complete genomes, I estimated that the *B. cereus s. l.* core and pan-genome contain ≈600 and ≈60,000 protein-coding genes, respectively. Pan-GWAS analysis revealed significant associations of particular genes with phenotypic traits shared by groups of taxa. All phylogenetic analyses recapitulated two previously used classification systems, and taxa were consistently assigned to the same major clade and group. By including accessory genes from the pan-genome in the phylogenetic analyses, I produced an exceptionally well-supported phylogeny of 114 complete *B. cereus s. l.* genomes. The best-performing methods were used to produce a phylogeny of all 498 publicly available *B. cereus s. l.* genomes, which was in turn used to compare three different classification systems and to test the monophyly status of various *B. cereus s. l.* species. The majority of the methodology used in this study is generic and could be leveraged to produce pan-genome estimates and similarly robust phylogenetic hypotheses for other bacterial groups.

## List of abbreviations

ACLAME: A CLAssification of Mobile genetic Elements
AFLP: amplified fragment length polymorphism
BAPS: Bayesian Analysis of Population Structure
BCSL: *Bacillus cereus sensu lato*
BLAST: Basic Local Alignment Search Tool
BP: bootstrap probability
GB: gigabytes
GWAS: genome-wide association study
HMM: hidden Markov model
HaMStR: Hidden Markov Model based Search for Orthologs using Reciprocity
LCB: locally collinear block
MAFFT: Multiple Alignment using Fast Fourier Transform
MGE: mobile genetic element
ML: maximum likelihood
MLST: multilocus sequence typing
MP: maximum parsimony
NCBI: National Center for Biotechnology Information
PANTHER: Protein ANalysis THrough Evolutionary Relationships
PATRIC: Pathosystems Resource Integration Center
PAUP*: Phylogenetic Analysis Using Parsimony *and other methods
PHYLIP: Phylogeny Inference Package
PRANK: Probabilistic Alignment Kit
RAM: random access memory
RAxML: Randomized Axelerated Maximum Likelihood
RF: Robinson-Foulds
RefSeq: Reference Sequence database
SNP: single nucleotide polymorphism
dDDH: digital DNA-DNA hybridization
gsi: genealogical sorting index
nt: nucleotides

## Declarations

### Acknowledgements

I thank Shashikala Ratnayake for assistance generating the Mash-distance-based phylogeny of *Bacillus*, Todd Treangen for helpful discussions about the project, and M.J. Rosovitz, Brian Janes, and Martina Eaton for providing feedback on drafts of the manuscript.

### Availability of data and material

All data analyzed during the current study were downloaded from public databases (ACLAME, NCBI, and PATRIC), and dates of download are provided in the text. A list of RefSeq assembly accessions for the taxa used in this study is provided in Additional file 1.

## Additional files

**Additional file 1 — RefSeq assembly accessions for the taxa used in this study.** A list of RefSeq assembly accessions for the bcsl_498 taxa.

**Additional file 2 — Roary gene presence/absence matrix for BCSL_114 taxa.** The gene presence/absence spreadsheet lists all genes in the pan-genome and the taxa in which they are present, along with summary statistics and additional information.

**Additional file 3 — Roary gene presence/absence matrix for BCSL_498 taxa.** The gene presence/absence spreadsheet lists all genes in the pan-genome and the taxa in which they are present, along with summary statistics and additional information.

**Additional file 4 — Binary matrix of phenotypic traits exhibited by BCSL_498 taxa.** Binary phenotypic trait matrix for bcsl_498 taxa, created using the isolate meta-data obtained from PATRIC.

**Additional file 5 — Construction of a HaMStR database.** Prokka was used to annotate 114 *B. cereus s. l.* complete genomes. The resulting protein-coding gene annotations were provided as input to Roary, which constructed a pan-genome consisting of 59,989 orthologous protein sequence clusters. After filtering, which included mobile genetic element removal, the 8,954 remaining clusters were aligned with MAFFT. Gene models were built from the multiple sequence alignments using the `hmmbuild` program from HMMER. The 8,954 gene models, together with separately constructed reference taxon BLAST databases, constituted the hamstr_full_mges_removed HaMStR database.

**Additional file 6 — Construction of a concatenated data matrix.** Prokka was used to annotate *B. cereus s. l.* “query genomes”— i.e., draft genomes that were not included in BCSL_114. The resulting protein-coding gene annotations were provided as input to HaMStR, which used the `hmmsearch` program from HMMER followed by BLASTP to assign query sequences to HaMStR database gene models. Clusters of orthologous protein sequences from query and database taxa were aligned with MAFFT and converted to corresponding nucleotide alignments. The multiple sequence alignments were reduced to a single sequence per taxon with a consensus procedure that used nucleotide ambiguity codes to combine information from sequence variants where necessary. The individual alignments were then concatenated to produce the final data matrix.

**Additional file 7 — Mash-distance-based phylogeny of the genus *Bacillus.*** Phylogeny of 146 *Bacillus* genomes, computed with Mash and FastME.

**Additional file 8 — Attributes of** *B. cereus s. l.* **species, clades, and groups.** Tables that provide number of taxa, average number of genes found among Roary clusters, and average number of genes present in mat_6 for *B. cereus s. l.* species, clades, and groups.

**Additional file 9 — Taxon metadata for bcsl_498.** Table providing clade, group and hierBAPS cluster affiliation for bcsl_498 taxa, along with the number of genes found among Roary clusters (complete genomes only) and the number of genes present in mat_6 (out of a possible total of 8,954 genes).

**Additional file 10 — Rarefaction curve: core vs. all genes.** The rarefaction curve shows that after ≈35 genomes have been sampled (≈31% of all genomes), the number of core genes remains fairly constant at ≈600 genes, while the total number of genes in the pan-genome continues to increase almost linearly.

**Additional file 11 — Rarefaction curve: new vs. unique genes.** The rarefaction curve shows that as genomes are sampled, genes never before observed continue to be found at a fairly steady rate, and the total number of unique genes discovered continues to increase, with no indication of soon approaching an asymptote.

**Additional file 12 — Accessory binary tree and gene presence/absence visualization.** The “accessory binary tree” and gene presence/absence information produced by Roary are plotted side-by-side. The outermost *B. cereus s. l.* clades include taxa with relatively few accessory genes included in the analysis, such as *B. cytotoxicus, B. mycoides*, and *B. pseudomycoides.* By contrast, the genomes with the most accessory genes present belong to the highly clonal clade of *B. anthracis* strains.

**Additional file 13 — Scoary result summary, including enriched gene ontology biological processes.** Positively or negatively trait-associated gene sets produced by Scoary were subsequently tested for possible enrichment of gene ontology biological processes. Complete Scoary results for eight traits, including gene annotations, are also given.

**Additional file 14 — Robinson-Foulds distance between all pairs of BCSL_114 phylogenetic results.** Both the standard and normalized Robinson-Foulds distance is given.

**Additional file 15 — BCSL_114 maximum likelihood phylogenetic analysis results.** Cladogram depicting the best estimate of the phylogenetic relationships among bcsl_114 taxa, computed with RAxML using 8,954 genes (ml_6; Table 4). *B. cytotoxicus* was used to root the tree. Major *B. cereus s. l.* clades and groups are indicated, as are bootstrap probabilities.

**Additional file 16 — BCSL_498 maximum likelihood phylogenetic analysis results, color-coded by species.** Phylogram depicting an estimate of the phylogenetic relationships among bcsl_498 taxa, computed with RAxML using 8,954 genes (ml_8; Table 4). *B. manliponensis* was used to root the tree. *B. cereus s. l.* species tested for monophyly with the *gsi* are color-coded.

